# Nest microbiota and pathogen abundance impact hatching success in sea turtle conservation

**DOI:** 10.1101/776773

**Authors:** Daphne Z Hoh, Yu-Fei Lin, Wei-An Liu, Siti Nordahliawate Mohamed Sidique, Isheng Jason Tsai

## Abstract

Hatchery practices are pivotal to conservation success. In sea turtle hatchery, reusing the same sand has been a norm but remains unclear whether such approach increases the risk of *Fusarium solani* species complex (FSSC) infection causing huge mortality in sea turtle eggs worldwide. We employed 16S and ITS amplicon sequencing in 123 sand samples and isolated fungal strains from diseased eggs across seven hatcheries and neighboring beaches in Malaysia. FSSC was isolated from all sampled hatcheries where *F. solani/falciforme* was the predominant species. A distinct microbial composition and higher abundance of FSSC (mean = 5.2 %) was found in all but one hatchery when compared to nesting beaches (mean = 1.3 %). Specifically, an ascomycetous fungus *Pseudallescheria boydii* consistently appeared in higher abundance (mean = 11.4 %) in FSSC-infected nests and was significantly associated with lower hatching success. The hatchery that maintained the most stringent practice by changing sand every nesting season had a microbiota resembling nesting beaches as well as lowest FSSC and *P. boydii* abundance. The results of current study imply the need to avoid reusing sand in sea turtle hatchery.

## Introduction

The establishment of hatchery has led to conservation success of endangered sea turtles. For instance, hatchery can reduce egg poaching, serve as a shelter under extreme weathers and facilitate conservation management[1, 2]. However, hatchery is an example of “all one’s eggs in one basket” where a single incidence can destroy all eggs[3]. Hence, strict hatchery practices have to be followed from the beginning since egg collection[2]. To maximize hatching success, extensive research efforts have been carried out on egg-handling procedures[1, 4] and incubation conditions[5–7]. Despite variability and constant revisions in these practices, a consensus has yet reached on choosing suitable substrates and environment for sea turtle egg incubation in a hatchery setting. Given the correct handling of eggs from nesting beach, establishing a new nest in a hatchery is a good measure to improve hatching success. Nonetheless, human interference on natural process such as egg relocation is usually discouraged[2]. One of the unforeseen anthropogenic induced impacts is the pathogen contamination of eggs in hatchery. Hatcheries were reported to have a higher disease incidence[8, 9] as high host density allowed pathogens to transmit more effectively[10]. Yet, little is known regarding disease occurrence on eggs in sea turtle hatchery in comparison with natural nesting beach.

*Fusarium falciforme* and *F. keratoplasticum*, part of the *Fusarium solani* species complex (FSSC)[11], are two pathogenic fungi known to infect sea turtle eggs with high mortality[12–14]. Koch postulates were fulfilled from infection assay using *F. keratoplasticum* on loggerhead turtle eggs[15], and this emerging fungal disease is now considered worldwide as more FSSC were isolated from dead sea turtle eggs of different continents in recent years[14, 16, 17]. Most infections occurred at natural nesting beaches[12–14] and few reported in hatcheries[17, 18]. Additionally, these fungi are known causative agents in other host species which includes human, animals and plants[19–22].

To improve disease management, an increasing number of studies had been carried out to investigate the relative abundance of pathogens within the microbiota to disease prevalence. Targeting a specific pathogen, these studies typically attempted to limit the abundances of such pathogens in the environment. Examples ranged from the characterization of microbial community after the use of suppressive soil in agricultural farms[23, 24] to post-probiotic treatment in an oyster hatchery[25]. Present studies on microbial community in sea turtle nesting beach focused only on *arribada* event, which primarily aimed to determine the association between microbial abundance or richness and hatching success[26, 27]. Since sand-borne origin of FSSC infection on eggs was confirmed[14], there is a need to encompass the relative abundances of pathogens for the analysis of microbial community structure in the environment for egg incubation. Past studies have successfully cultured FSSC from the nest sand and debris from nest surroundings[16, 17]. However, the ubiquitous nature of FSSC[19] implies that successful infection depends on the extent of natural nests experiencing environmental stressors such as tidal inundation and unsuitable substrate’s sizes[14]. One unintended anthropogenic stressor may be the re-establishment of nests in hatcheries, and little is known how this contributes to disease occurrence.

In Peninsular Malaysia, eggs laid at the nesting beach are relocated to the nearest *ex-situ* hatchery to reduce poaching[28, 29]. However, some hatcheries have recently experienced FSSC infection[17]. Most *ex-situ* hatcheries in Peninsular Malaysia reuse the same sand within the hatchery (a fenced area) for egg incubation for several nesting seasons. This approach had long been discouraged with the suspicion that sand containing egg residues from previous seasons may act as a reservoir for pathogens or toxic byproducts[3]. We hypothesized that microbial community and FSSC abundance in the reused-sand of hatchery differed with natural nesting beach. In this study, we investigated if FSSC could be successfully identified from diseased eggs across sea turtle hatcheries of Peninsular Malaysia. We profiled the microbial composition of sands collected in hatcheries and their residing nesting beaches. Accordingly, we contrasted the differences between two environments and identified species associated with hatchery sands. We further examined whether their abundances were associated with hatching success. Together, the study discloses microbiome changes contributed by hatchery practices and offers new insights to inform conservation practices in sea turtles.

## Materials and methods

### Site description and sampling

Sampling permissions were obtained through verbal consent from all sites. We collected samples from seven sea turtle hatcheries across Peninsular Malaysia in July 2018. Descriptions of these sites are shown in Fig. 1. Nest density and clutch size of each site were estimated based on available reports (Table S1). Sampled nests from all hatcheries were of *Chelonia mydas* except nests from Melaka (M) site were *Eretmochelys imbricata*. A total of 123 sand samples were collected from hatcheries’ nests which were used to incubate eggs (henceforth referred to as hatchery sand) and the nearest turtle nesting beach to each hatchery (control sand). Hatchery sands were collected within two days from nests where hatchlings have emerged. The only exception was except Geliga (G) where permission was given to collect sands from total failed nests only. Control sand was collected approximately 60 cm deep from the beach surface to mimic natural nesting settings of sea turtles and avoided areas with vegetation. Two to three such regions were chosen from each nesting beach. Three technical replicate samples were collected from each nest in hatchery or region in nesting beach. All sand samples were stored at – 20 °C until DNA extraction. Hatching success was calculated averaging three excavated nests in each site (Fig. 1). All samples were brought into Taiwan with the permission from Bureau of Animal and Plant Health Inspection and Quarantine under permit No. 107-F-501.

**Figure 1:**
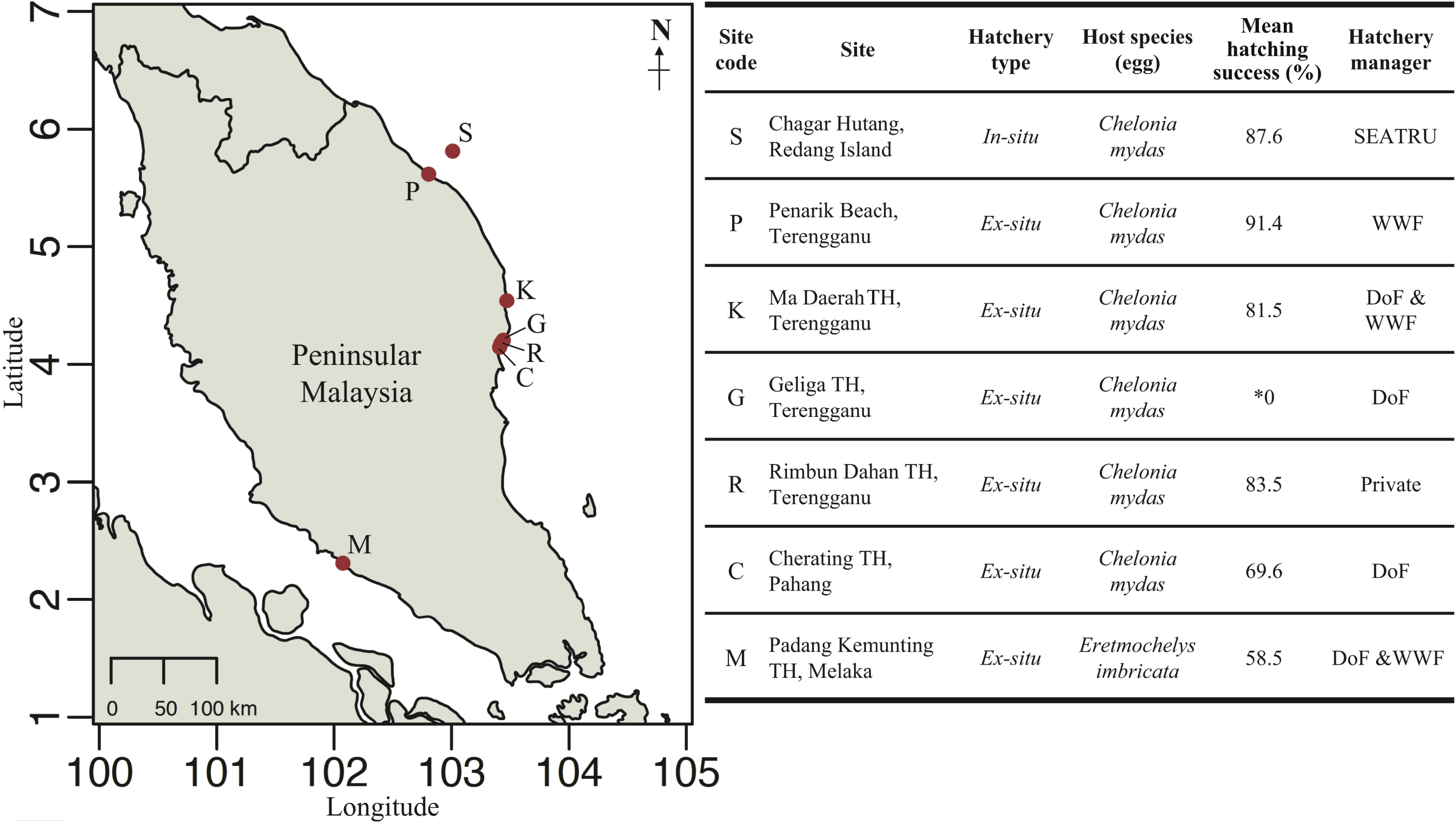
Map, location and metadata of the sampled sea turtle hatcheries and neighboring beaches in Peninsular Malaysia. TH, turtle hatchery; SEATRU, Sea Turtle Research Unit, Universiti Malaysia Terengganu; WWF, World Wide Fund for Nature, Malaysia; DoF, Department of Fisheries Malaysia. *Asterisk denotes that samples were collected from total failed nest only.

### *Fusarium* isolates from diseased eggs

For each excavated nest, we randomly collected three *Fusarium*-infected dead eggs based on symptoms as described previously[15] for isolation of *Fusarium* species. Eggshells were rinsed with distilled water to remove sand particles, cut into approximately 1 × 1 cm using sterilized scissors and incubated on Potato Dextrose Agar with 50 μg/mL chloramphenicol at 30 °C. Subcultures were made on the same medium to obtain pure culture. The primer pairs SR6R/ITS4[30] was used to run PCR amplification of the internal transcribed region of these isolates. Phylogenetic analysis was performed with representative isolates from each site and nest along with selected *Fusarium* sequences obtained from NCBI nucleotide database. Sequences were trimmed using trimAl with ‘automated1’ option (v1.2rev59)[31], aligned using MAFFT for 1 000 iterations (v7.310)[32], and constructed phylogenetic tree using FastTree with nucleotide alignment assuming generalized time-reversible model (v2.1.10)[33]. Tree visualization was made with FigTree software (v1.4.4)[34].

### DNA extraction and amplicon library generation for high-throughput sequencing

Total genomic DNA was extracted from sand samples using DNeasy PowerSoil Kit (QIAGEN, Hilden, Germany, Cat. #12888) with minor modifications to the manufacturer’s instructions. One gram of homogenized sand was used in each extraction. PowerLyzer 24 homogenizer (MoBio Laboratories, Carlsbad, CA, USA, Cat. #13155) was set to 3 000 rpm for 10 min. DNA was eluted with 60 μl of Solution C6. Bacterial 16S rRNA and fungal internal transcribed spacer (ITS) amplicons were generated using tagged primer pairs V3/V4[35] and ITS3ngs(mix)/ITS4[36] respectively. PCR reaction and thermocycling condition are as previously described[36]. Positive and no-template controls (elution buffer) were included in the PCR reaction and amplicon pool. ZymoBIOMIC Microbial Community DNA Standard (Zymo Research Corporation, Irvine, CA, USA, Cat. #D6305) was used as the mock community control in 16S amplicon preparation. For ITS amplicon, equal DNA concentration of *Fusarium solani, Mycena venus*, and *Mortierella elongata* was used as single organism control. Amplicons were normalized with SequelPrep Normalization Plate (96) Kit (Invitrogen Corporation, Carlsbad, CA, USA, Cat. #A10510-01). Normalized amplicons were pooled at equal volume and concentrated using Agencourt AMPure XP beads (Beckman Coulter, Brea, CA, USA, Cat. #A63881). ITS amplicon library was prepared using Illumina TruSeq DNA Prep Kit. Sequencing was performed by the NGS High Throughput Genomics Core in Biodiversity Research Centre, Academia Sinica, Taiwan. All amplicon libraries were sequenced using Illumina MiSeq with paired-end 2 × 300 bp chemistry.

### Sequence data processing

Raw fastq sequences were demultiplexed according to their respective barcode using sabre (v1.0; https://github.com/najoshi/sabre) allowing for one nucleotide mismatch. Adaptors and primer sequences were trimmed using USEARCH (v10)[37]. Processing of sequence reads was carried out following the UPARSE pipeline[38]. In brief, forward and reverse reads were merged and filtered using the default settings in USEARCH. Operational taxonomic unit (OTU) clustering at 97 % sequence and minimum OTU abundance was set to 8 during the OTU clustering to denoise the data and remove singletons. OTU table was created using *usearch_global* option. Taxonomy classification of OTU was performed against RDP training set (v16) and UNITE database (v7.1) for 16S and ITS reads respectively using the SINTAX algorithm[39, 40].

### *Fusarium solani* species complex (FSSC) species-level analyses

Due to incomplete taxonomy assignment through the UNITE database, 38 OTU sequences assigned as Nectriaceae were manually re-identified by comparing against NCBI nucleotide database using BLAST. The OTUs which are *Fusarium* genus according to blast result were renamed in the *tax_table* of *phyloseq* object before further analyses. Blast result from four out of the 38 OTUs were three *Fusarium solani* and a *Fusarium keratoplasticum*. These four OTUs were then grouped as a single OTU being part of the FSSC[41] and known to infect sea turtle eggs[14, 42].

### Microbiota community analyses

Prior to analysis, the validity of sequence data was confirmed by comparing the known identity of positive controls against the taxonomic classification result from UPARSE pipeline. Negative control ensured the absence of contamination in the samples. Technical replicates were merged having found no significant variation in their microbial compositions. Unclassified ITS OTU sequences were manually curated by comparing against NCBI nucleotide database using BLAST. Non-fungal eukaryotic OTUs were then removed from further analyses. The bulk of the analyses were carried out using R-Studio (v1.1.463)[43]. Amplicon data were analyzed with *phyloseq* package (v1.28.0)[44]. For beta diversity analysis, OTUs that appeared less than three times in at least 20 % of all samples were filtered from the analysis. Distances between samples were calculated using Bray-Curtis dissimilarity. Heatmap was plotted using *pheatmap* package (v1.0.12)[45] with distance measure using “correlation” option to perform samples clustering. Differentially abundant taxa were determined using DESeq2 (v1.24)[46] at genus and OTU level for bacterial and fungal amplicons, respectively. Sample filtering criteria for differentially abundant taxa were as per in beta diversity analysis with the addition where hatchery sand samples from site P were omitted.

Having potential biomarkers determined through heatmap and DESeq2 analyses, we further constructed correlation networks to determine their relationship. Fastspar (v0.0.9)[47], an implementation of Sparse Correlations for Compositional Data[48] algorithm was used to estimate correlations between each bacterial genus and fungal OTUs from the combined 16S and ITS compositional data. The output matrices were filtered to keep correlation value > 0.3 and with p-value < 0.05. Networks were visualized using Cytoscape (v3.7.2)[49].

## Results

### Data characteristics

To determine the effect of reusing sand on microbiota structure and pathogens’ occurrence and abundance, we sampled 63 *Fusarium*-infected dead eggs and 123 sand samples (63 hatchery sands and 60 control sands) from seven sea turtle hatcheries across Peninsular Malaysia (Fig. 1). All sampled hatcheries were *ex-situ* except Chagar Hutang (Site code: S), which is the only hatchery with an *in-situ* practice. Among the hatcheries, Penarik (P) relocates the whole hatchery to a new location after every nesting season, which makes it the only hatchery that does not reuse the same sand for egg incubation. All sampled nests contained dead eggs with symptoms of fungal colonization as previously described[15, 50]. Upon molecular characterization of fungi isolated, *Fusarium solani* species complex (FSSC) were identified from all hatcheries (Fig. S1). Of these, 70 (84.3 %) were *F. solani/falciforme*, 12 (14.5 %) *F. keratoplasticum*, and one *F. oxysporum* isolate.

### Distinct microbial community in FSSC-infected sea turtle nest

Overall, sand microbiota in sea turtle nesting beaches and hatcheries were composed of 7 269 bacterial and 3 287 fungal OTUs. Phyla compositions in bacterial and fungal were similar regardless of sand types (Fig. S2). The most dominating bacterial phylum across all samples was *Proteobacteria* (averaging 36.6 %), which was followed by unclassified bacteria (15.4 %) and *Firmicutes* (13.3 %) (Fig. S2A). Between sand types, most of the bacterial phyla were significantly different in relative abundance (Fig. S3A). In particular, *Bacteroidetes* was almost twice in abundances in hatcheries (13.3 %) than nesting beaches (5.7 %). This was, however, due to an elevated relative abundance of over 20 % in four nests from three hatcheries (MN2, MN3, SN1, and RN1; Fig. S2A) suggesting that large variations exist between and within the same site. No single *Bacteroidetes* OTU was responsible for the increased proportion. Bacterial species richness and community evenness indices were significantly lower in hatchery sand compared to nesting beach (Fig. S4A). In the case of fungi, the phylum Ascomycota has the highest mean relative abundance (38.1 %), consistent with the frequent observation of dominant fungal species present in soil communities[51], followed by unclassified fungi (35.9 %) and Basidiomycota (22.3 %) (Fig. S2B). Four nests and three beach samples in hatchery G and P have shown higher Basidiomycota than Ascomycota relative abundance due to the dominance of *Inocybe* sp. ranging from 49.3 % to 97.5 %. Mean relative abundances of four major fungal phyla were similar between sand types (Fig. S3B). In contrast to the bacterial community, hatchery sand harbored an increased fungal richness but reduced community evenness (Fig. S4B).

Microbial communities of sand samples were significantly different between hatcheries and nesting beaches (Fig. 2, Fig. S5). Compared to five *ex-situ* hatcheries that reused sand, P hatchery stood out as the only hatchery changing location every nesting season and have exhibited similar bacterial community to nesting beaches (Fig. S5). Statistical tests showed significant difference and dispersion in bacterial composition between hatcheries and nesting beaches (after removing P hatchery sands; ANOSIM R = 0.96, *P* = 0.001; PERMANOVA R^2^ = 0.71, *P* < 0.001; betadisper *P* = 0.017). In the case of fungal community, samples consisted high proportion of *Inocybe* sp. were separated from beach and hatchery sand clusters (Fig. 2B). PC1 (21.3 %) set apart these samples to the rest while PC2 separated hatcheries and nesting beaches. After excluding these samples, similar conclusion was also drawn from the fungal community that significant dispersion and composition was observed between sand types (after removing samples with high *Inocybe* sp. relative abundance; ANOSIM R = 0.47, *P* = 0.001; PERMANOVA R^2^ = 0.21, *P* < 0.001; betadisper *P* = 0.001). Additionally, both bacterial and fungal community structures showed no geographical difference in either sand type (excluding P hatchery sands; In nesting beach or hatchery, 16S: *P* = 0.3 or 0.9; ITS: *P* = 0.4 or 0.9).

**Figure 2:**
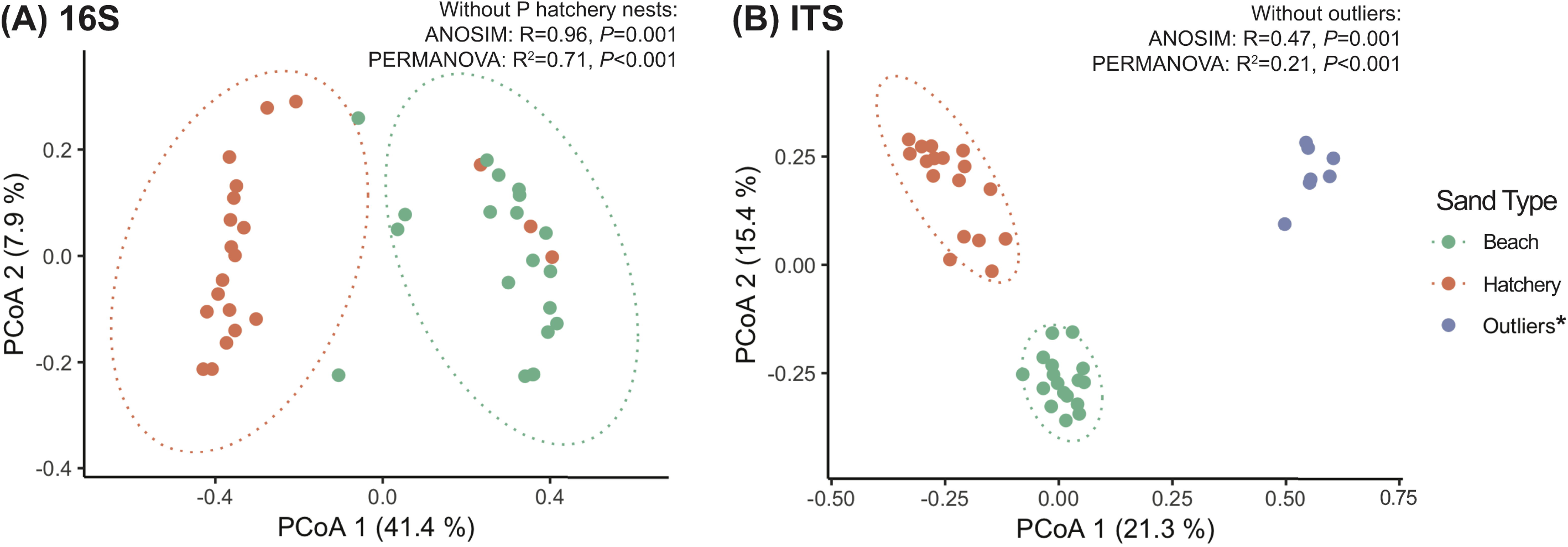
Principal Coordinate Analysis plots measured by Bray-Curtis distance of (A) bacterial and (B) fungal communities. Each dot represents one site and different color denote sand type. Ellipses represent 95 % confidence interval within each category (hatchery or nesting beach). Multivariate statistical analyses ANOSIM and PERMANOVA were used to test the community differences between sand types. *Asterisk denote a subset of samples containing high relative abundance of *Inocybe* sp.

### Differentially abundant taxa in FSSC-infected sea turtle nest

Clustering of relative abundances of top ten major taxa have partitioned the samples based on sand types, except P hatchery grouped with nesting beaches (Fig. 3). FSSC, *Pseudallescheria boydii* and an unclassified fungus (OTU5) were present with higher abundances in most hatcheries except P hatchery. The closest match of this unclassified fungal OTU5 to the public database belonged to the order *Capnodiales*, which contains fungal species that are plant pathogens[52]. *Saccharomyces cerevisiae* was more frequently detected in nesting beaches of S and M regions. Interestingly, nesting beach S had a relatively higher abundance of various *Bacillus* spp. (mean 9.7 % vs. averaging 7.2 % in all samples) and FSSC (3.0 % vs. averaging 1.4 % in all samples) when compared to other nesting beaches.

**Figure 3:**
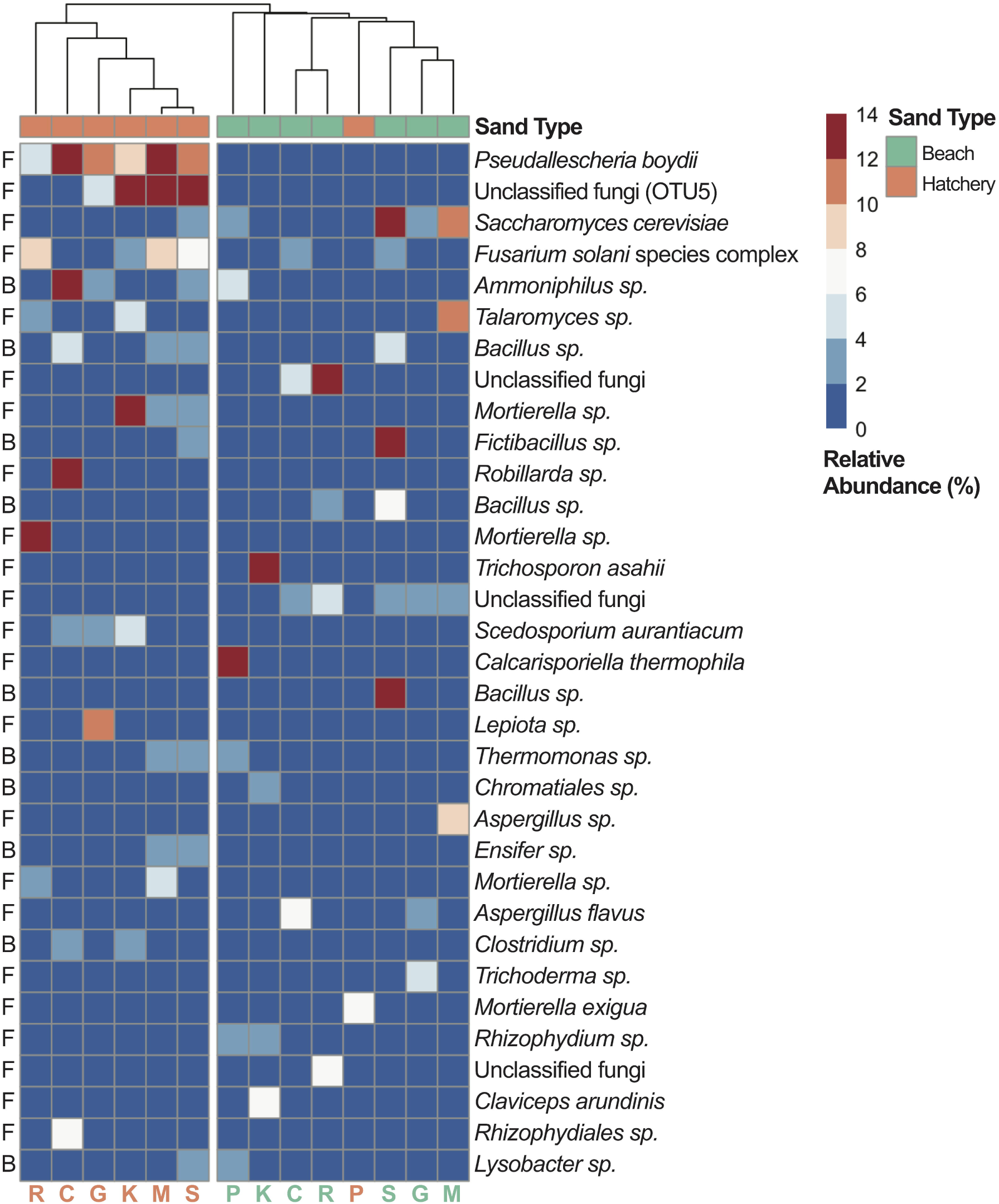
Heatmap of relative abundance (%) of top ten major bacterial and fungal OTUs from each site totaling 33 OTUs. Color in each box denote the mean relative abundance of samples in corresponding sites. F, fungal; B, bacterial.

Analysis of differentially abundant taxa revealed 106 bacterial genera and 48 fungal OTUs significantly higher in hatchery sand (Fig. S6 & S7). Conversely, nesting beaches harbored 45 and 94 higher abundance of bacterial genera and fungal OTUs, respectively. The aforementioned *P. boydii* and *Capnodiales* sp. were indeed significantly more abundant in the hatcheries (*P* = 5.36e^-15^ and 6.61e^-11^, respectively). Interestingly, FSSC was not significantly more abundant in hatcheries (*P =* 0.55). Upon closer inspection, FSSC also exhibited relatively high proportions in the nesting beaches of site S and C, implying the abundance of this fungal group can be high in nature. We next sought if there was an association between hatching success and summed abundances of differentially more abundant OTUs in nests. A significant negative correlation was found in fungal OTUs (included FSSC; Spearman correlation, ρ = - 0.59, *P* = 0.01; Fig. 4), and a negative but insignificant trend was observed in bacterial genera (Spearman correlation, ρ = - 0.29, *P* = 0.25). The differential bacterial and fungal OTUs had contributed a total increased abundance of averaging 32.9 % and 47.8 % in hatchery sand respectively, suggesting a higher occurrence of these OTUs coincide with the infection and may contribute towards a lower hatching success. Further independent analysis supported the differential abundances of taxa in different sand types. For instance, multivariate analysis also showed a higher abundance contribution of FSSC and *P. boydii* in nests (Fig. S8).

**Figure 4:**
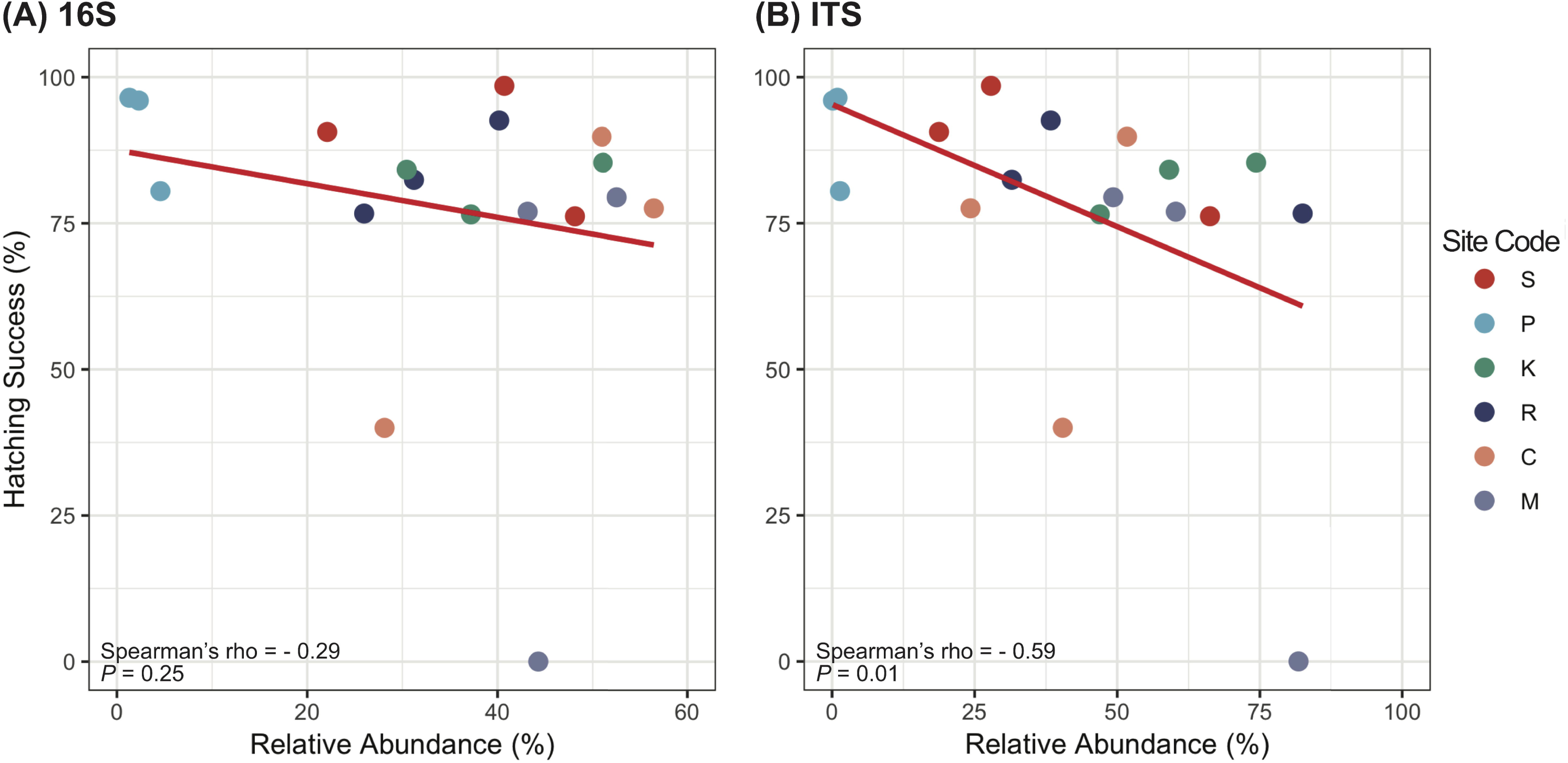
Scatterplots showing correlation between summed relative abundance of differentially abundant (A) bacterial genera and (B) fungal OTUs in sand and hatching success of each nest. Each dot represents one nest. Samples originated from G hatchery were excluded from statistical analysis due to nests consist solely of failed eggs, i.e., 0 % hatching success.

### Higher abundance of FSSC and *P. boydii* in reused-sand

We detected a higher mean relative abundance of FSSC in hatcheries compared with nesting beaches (excluded P hatchery; nesting beach vs. hatchery, 1.3 % vs. 5.2 %, *P* = 0.0017; Fig. 5). Of all the *Fusarium* species detected, FSSC comprised 94.9 % of all *Fusarium* sequences in the hatcheries. *P. boydii* also showed higher mean relative abundance in hatcheries (excluded P hatchery; unpaired-Wilcoxon test, nesting beach vs. hatchery, 0.3 % vs. 11.4 %, *P* = 2.5e^-07^; Fig. 5). *P. boydii* was found to be more consistently high in abundance across hatcheries (at least > 5 %) and on average 11.9 % higher than FSSC. There was no correlation between hatching success and FSSC’s relative abundance (Spearman correlation, ρ = - 0.1, *P* = 0.69). In contrast, a negative correlation was observed when compared to relative abundance of *P. boydii* (Spearman correlation, ρ = - 0.48, *P* = 0.04; Fig. 6). To assess whether this was not due to data dependence and true interaction exist between these two species[48], we constructed co-occurrence networks using combined compositional data of bacterial and fungal species (Fig. S9). There was no direct interaction between FSSC and *P. boydii* in the sand environment of either nesting beach or hatchery.

**Figure 5:**
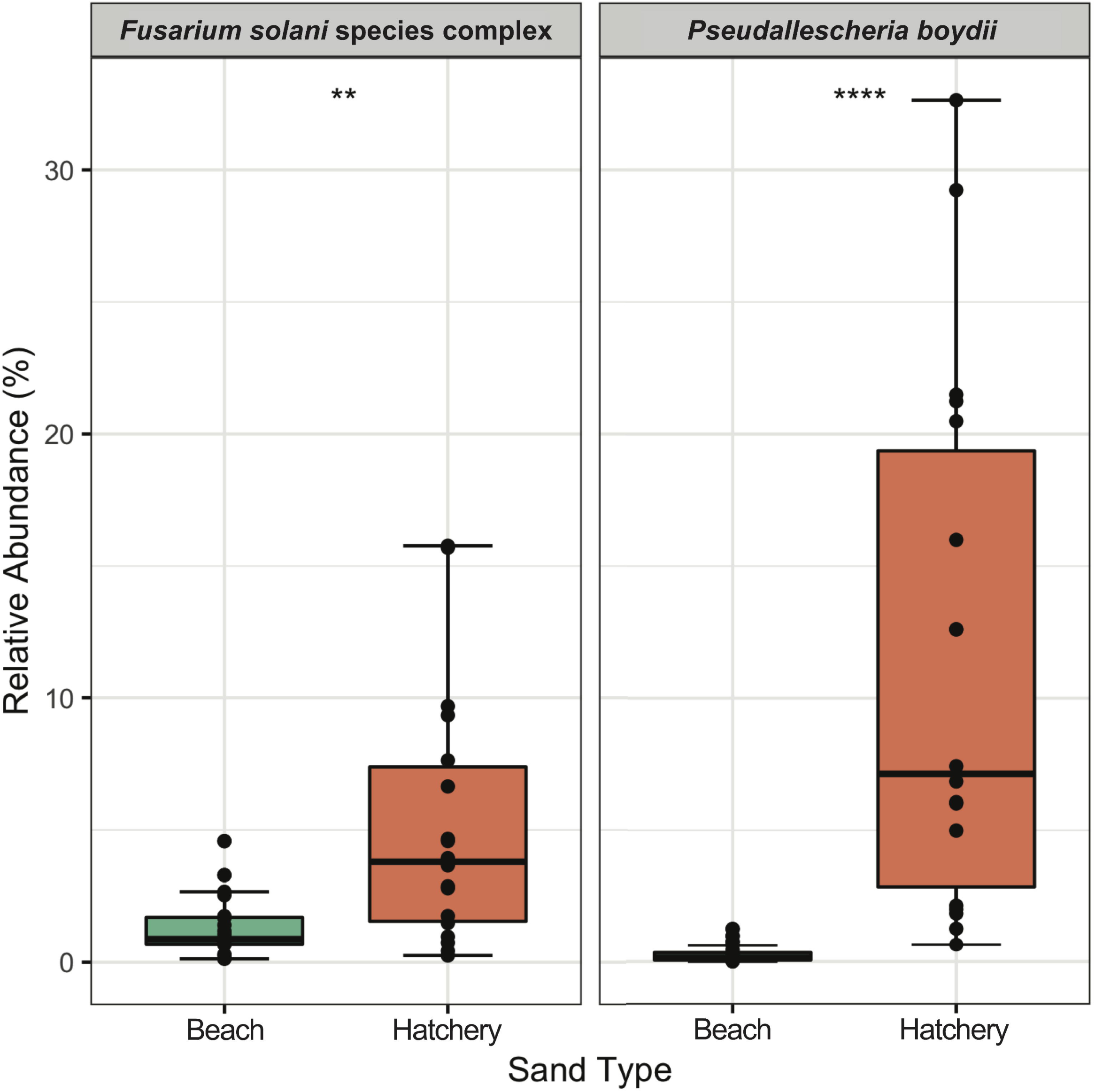
Boxplots showing relative abundance (%) of *Fusarium solani* species complex and *Pseudallescheria boydii* between two sand types. Statistical significance was calculated using unpaired-Wilcoxon test (** *P* ≤ 0.01; **** *P* ≤ 0.0001). Samples from P hatchery were removed from analysis due to evidently different community structures with other hatchery sand samples.

**Figure 6:**
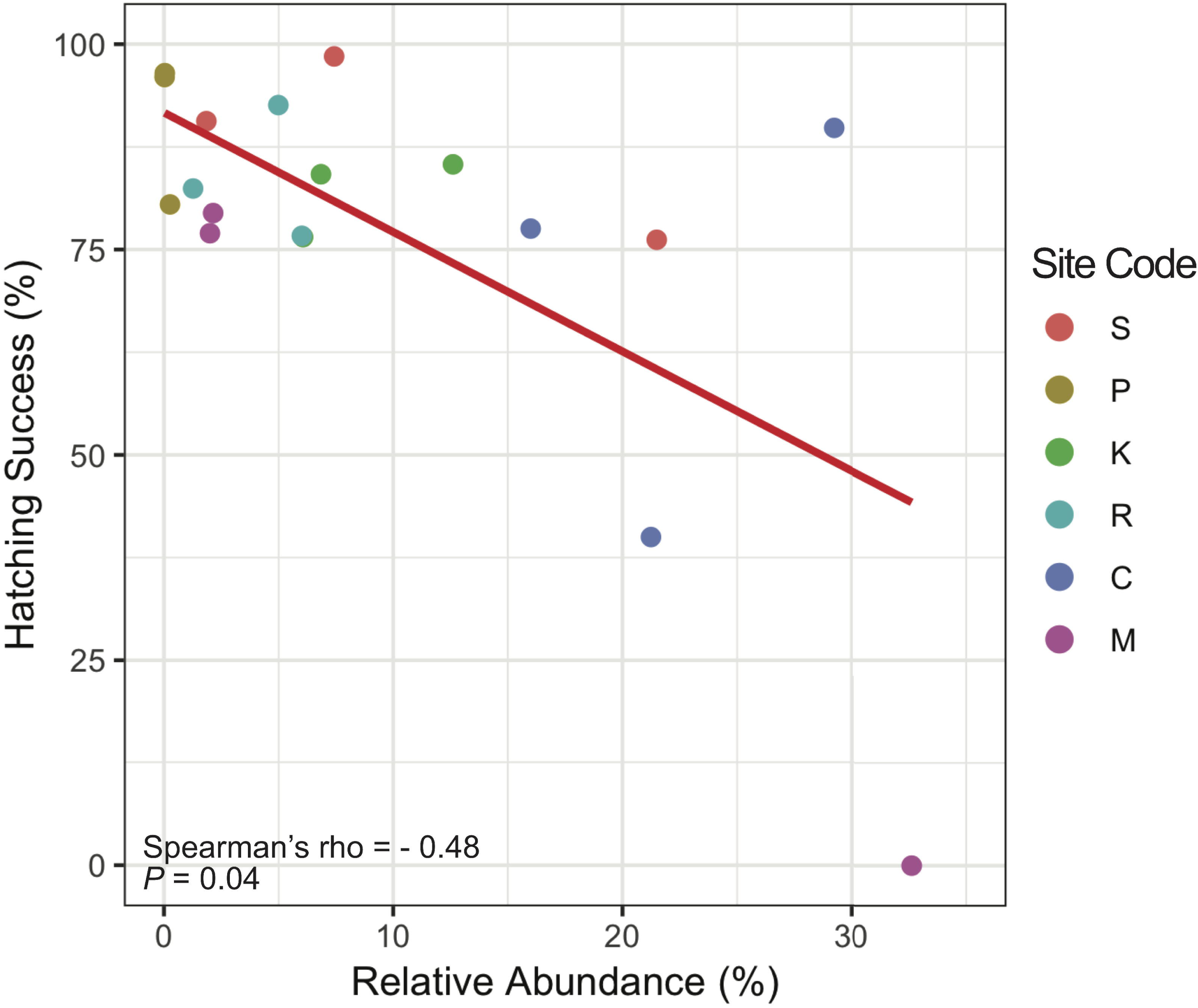
Scatterplot showing correlation between relative abundance of *Pseudallescheria boydii* in hatchery sand and hatching success of each sampled nests. Each dot is one nest. Samples of G hatchery sands were removed from analysis due to sampling of nests with all failed eggs only.

## Discussion

Previous reports have shown that FSSC present in natural nesting beaches can infect sea turtle eggs resulting in mass mortalities[14, 53]. To determine if different hatchery practices have an effect on disease prevalence, we determined the relative differences in FSSC and microbial composition of sand across multiple sea turtle hatcheries and neighboring nesting beaches in Peninsular Malaysia. Our findings revealed that all hatcheries that practiced sand reusage shared a distinctive microbiota that clearly differentiated from those of neighboring nesting beaches and hatchery P. By not reusing sand, hatchery P also harbored a lower relative abundance of FSSC among hatchery nests and had higher hatching success. We also qualitatively observed fewer eggs exhibited fungal-infected symptoms in this hatchery. The contrasting observations as a result of hatchery practices resolved the long speculation that reused sands indeed had a distinctive microbiota with higher relative abundances of FSSC. In addition, high abundances of several *Bacillus* spp. and FSSC were observed in the nesting beaches of the only *in-situ* hatchery S sampled in this study (Fig. 3). One possible explanation was due to high nesting turnouts which alter microbiota more drastically in a short nesting beach (approximately 350 m) with high nesting density (mean 1150 nests per year)[54].

The enclosed hatchery area may be a nutrient-rich cultivation environment of potential pathogens. Similar scenarios have been observed for other fungal diseases in animals and plants. The fungal pathogen of bats, *Geomyces destructans* was detected in the soil of ground environments where the bats hibernate[55]. Crop’s pathogens were also detected in the crop residues post-harvest, which can serve as sources of inoculum in the following season[56]. Reusage of sand will thus act a natural reservoir allowing FSSC to accumulate and infect eggs in subsequent nesting seasons. Our estimate of fungal relative abundance was only determined post-hatchling emergence and nest excavation to avoid eggs disturbance so that hatching success could be measured. Hence, the relative abundance of FSSC in hatcheries was a result of infection *a posteriori* and was not correlated to hatching success. Experiments had suggested colonization and the spread of fungal hyphae from diseased eggs to adjacent viable eggs[50], suggesting initial establishment is key to successful infection. Initial fungal density may be reduced from suitable sand treatments to increase hatching success[27] but may be infeasible due to space, cost and labor constraint.

FSSC is commonly found in the environment[19], along with other *Fusarium* species in natural nesting beaches. We were able to detect FSSC in higher abundance across hatcheries and in some of the nesting beaches, suggesting additional factors may contribute to the overall accumulation of this species complex. Differences in geographical distribution of *F. solani/falciforme* and *F. keratoplasticum* were observed. Previous reports in some of the main sea turtle nesting beaches in the Indian and Atlantic Oceans have shown that *F. keratoplasticum* predominated[14, 57], but instead we and recent survey[17] have more frequently isolated *F. falciforme* in Malaysia. Our study further revealed different abundances in FSSC species within Peninsular Malaysia. Among the hatcheries surveyed, only Melaka (M) hatchery was found with a higher relative abundance of *F. keratoplasticum* (∼ 5 %) with an increased isolation frequency[17]. Source of *F. keratoplasticum* in the environments have been associated with human activities[19]. However, it is not known whether such reason was the source as all sampled hatcheries have restricted access.

Various studies had shown that both FSSC and *P. boydii* are frequently isolated from fungal-infected sea turtle eggs[50, 57, 58]. Co-occurrence of the two fungi and the even higher prevalence of *P. boydii* suggest that it can be a more sensitive biomarker for FSSC-contaminated nest. *P. boydii* was found associated with lower hatching success, which warrants further examination on its status as a potential pathogen. To date, *P. boydii* was not proven to infect sea turtle eggs but the possibility exists since its ability to infect veterinary animals [59]. Hence, the high occurrence of *P. boydii* may be the consequences of the post-effect by FSSC infection. Interestingly, our co-occurrence network analyses suggest these two species did not directly interact with each other. Since *P. boydii* is a common soil microbe[60] and an opportunist pathogen which caused chronic diseases in human[60, 61], we speculate that FSSC may infect turtle eggs and *P. boydii* proliferate in masses by obtaining nutrients from dead eggs. In addition, lack of interaction between these two species also suggests FSSC alone can cause egg infection, which is consistent with the inoculation experiment[15]. Moreover, it is worth noting that a member of *P. boydii* species complex, *Scedosporium aurantiacum*[60] was found in hatchery sands in high abundance (see Fig. 3 & Fig. S7). This fungus was also previously isolated from hatchery sand[16] and diseased eggs[57], which provided evidence that our inferred interactions were accurate.

We have shown that the relative abundance of differentially abundant fungal species in hatchery sand were negatively correlated to hatching success. In a previous study, higher fungal abundance associated with lower hatching success had been demonstrated in nest sand of *arribada* beach[27]. The low hatching success was the ramification of the high microbial activity causing higher temperature and oxygen deprivation in nests[27, 62]. The hatchery nest environment is analogous to *arribada*. However, whether similar conditions are happening in hatchery warrant further study. Although no bacterial genera were associated with hatching success, we detected pathogenic bacteria such as the *Salmonella* and *Pseudomonas* spp. were significantly more abundant in hatchery sand. These species were previously isolated from failed sea turtle eggs[63] and recreational beaches[64, 65], potentially increase the infection risk to both hatchery personnel and eggs.

To the best of our knowledge, this study is the first to delineate the microbiota and quantify FSSC abundance in the nests of sea turtle hatcheries. Our key findings indicated that FSSC abundance and infection was most probably heightened from the practice of reusing sand for egg incubation. The consequence of such result suggests that turtle eggs and hatchery staffs suffer an increased risk of the infection by FSSC or other opportunistic microbes. Hence, we recommend sea turtle hatchery to change sand after every nesting season in order to maximize hatchling productions. This work emphasizes that stringency in hatchery management must be maintained for the efforts of conservation to not be in vain. For disease control and prevention, future attention needs to be paid to on the causes and consequences of abundances and interactions of microbes at different stages of infection not just in nests but also the nesting beaches due to rapid changing environments.

## Acknowledgements

We thank Department of Fisheries Malaysia, World Wide Fund for Nature Malaysia, Sea Turtle Research Unit, Universiti Malaysia Terengganu, and Rimbun Dahan Turtle Hatchery for the permission in sample collection. We thank Dr. Muhammad Hafiz Borkhanuddin, Ms. Noorkhalilie bt Che Abd Aziz, and Mr. Hsueh-yu Lu for the sampling arrangement and assistance. We also thank Ms. Chian-Mei Wei, Ms. Yi-Hua Chen, and Ms. Jeng-Yi Li in High Throughput Genomics Core at Biodiversity Research Center, Academia Sinica for sequencing. We thank Chia-Lin Chung for critically commenting on the manuscript.

## Author Contribution

IJT conceived the study. DZH, YFL, SNS and IJT designed the study. DZH and SNS performed the sampling. DZH, YFL, and WAL carried out experiments. DZH and YFL analyzed the data. DZH, YFL and IJT wrote the manuscript. All authors read and approved the final manuscript.

## Data Availability

Supplementary information is available at ISMEJ’s website. Sequences of isolates were deposited at GenBank under accession numbers MN326633-63. Amplicon sequence data were deposited in NCBI Sequence Read Archive under BioProject PRJNA560331.

