## Supplementary Information for "Nest microbiota and pathogen abundance impact hatching success in sea turtle conservation"

This file includes Table S1 and Figure S1 to S9.

**Table S1:** Site information

| Site code | Site | Mean number of nests per season | Mean number of eggs per nest | References |
|---|---|---|---|---|
| S | Chagar Hutang, Redang Island | 1500 | NA | [1] |
| P | Penarik Beach, Terengganu | 122 | 75 | [2] |
| K | Ma Daerah TH, Terengganu | 128 | 84 | [2] |
| G | Geliga TH, Terengganu | 924 | 95 | [2] |
| R | Rimbun Dahan TH, Terengganu | NA | NA | NA |
| C | Cherating TH, Pahang | NA | NA | NA |
| M | Padang Kemunting TH, Melaka | 416 | 113 | [3, 4] |

TH, turtle hatchery

[1] Chan EH. A 16-year record of green and hawksbill turtle nesting activity at Chagar Hutang Turtle Sanctuary, Redang Island, Malaysia. Indian Ocean Turtle Newsletter. 2010; **12**: 1–5.

[2] Department of Fisheries Malaysia of Terengganu. Report of sea turtle conservation center of Rantau Abang. Department of Fisheries Malaysia of Terengganu: Rantau Abang; 2017. (In Malay).

[3] Farisha NZA. Marine: Towards a better sea turtle awareness and conservation across Malaysia. *WWF-Malaysia Annual Review*. WWF-Malaysia: Petaling Jaya, Selangor, Malaysia, 2016, pp 14.

[4] Wong S. Marine: Notable marine turtle quantitative figures. *WWF-Malaysia Annual Review, 45 Years of Conservation Action.* WWF-Malaysia: Petaling Jaya, Selangor, Malaysia, 2017, pp 9.

**Figure S1:** Phylogenetic tree of *Fusarium* strains isolated from diseased eggs. Only selected *Fusarium solani* species complex isolates from this study were shown. Numbers at node indicate bootstrap percentages of clade supported out of 1000 replicates.

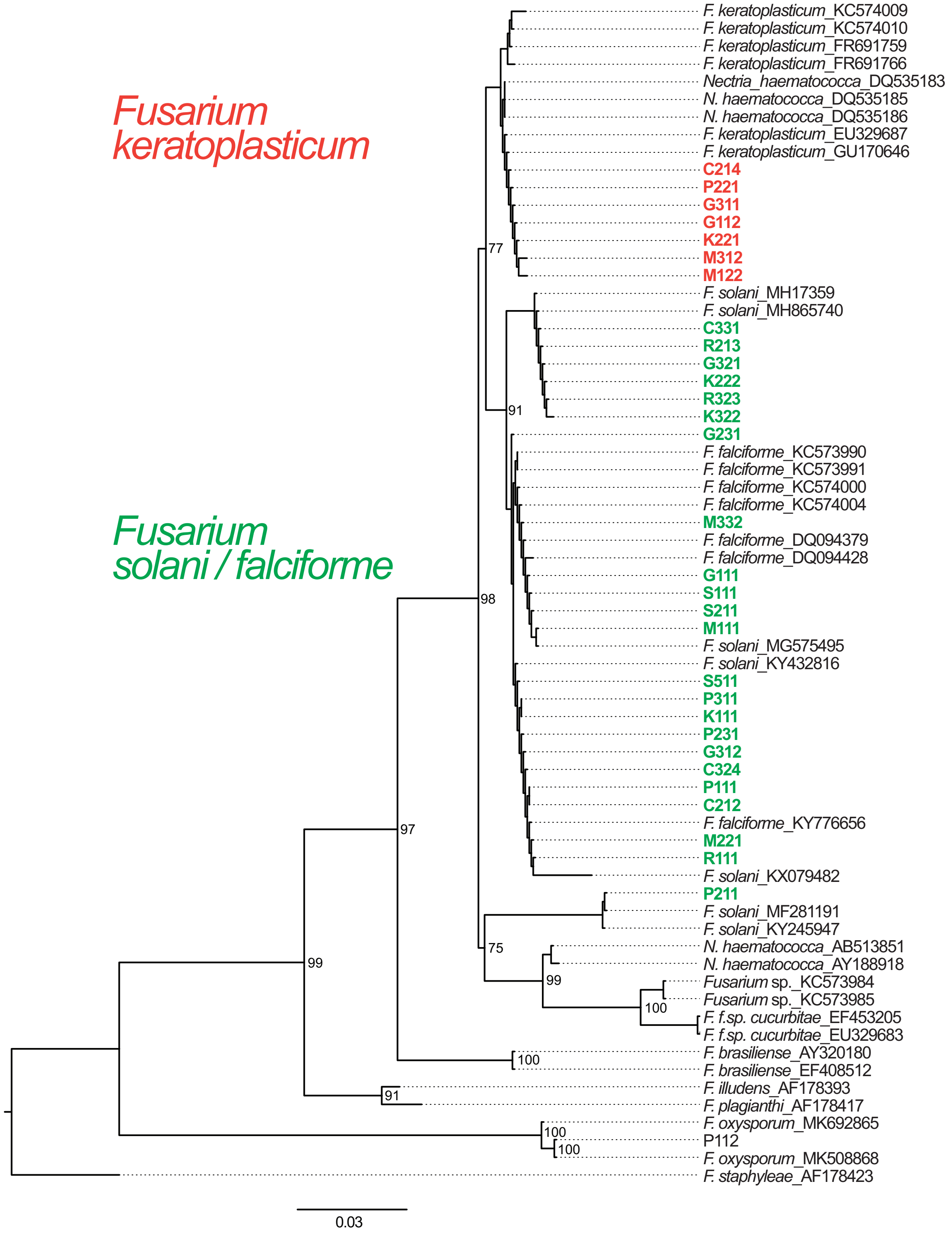

**Figure S2:** Bar plots of bacterial (A) and fungal (B) relative composition in all samples at the phylum level. First and second alphabet in Sample ID represents site code and sand type (B: Beach; H: Hatchery), respectively. *Asterisks indicate samples contained high proportion of *Inocybe* sp.

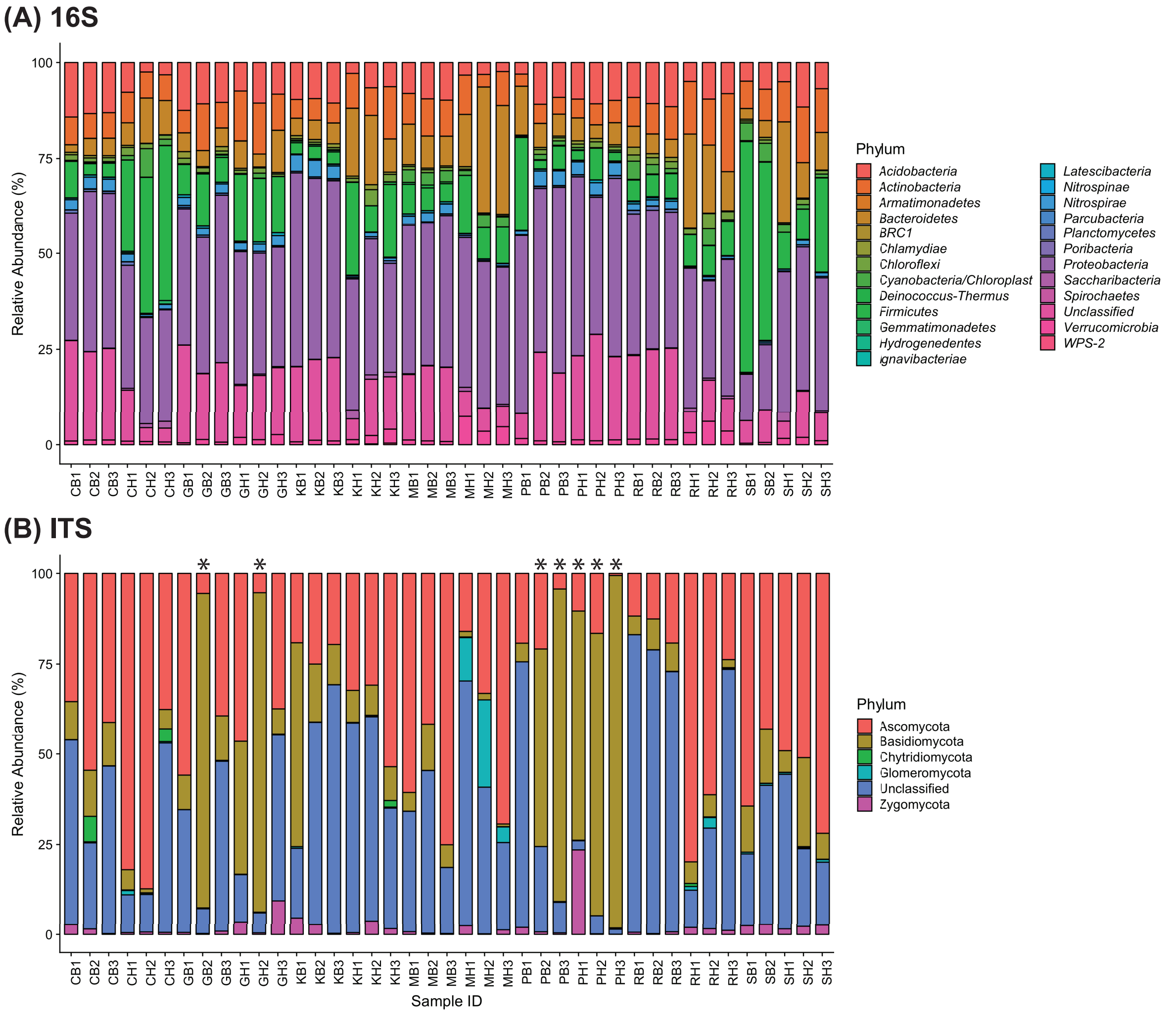

**Figure S3:** Box plots of bacterial (A) and fungal (B) relative composition of different sand types at phylum level. Each dot represents one sample. Only the top eight major bacterial phyla are shown in (A). Statistical significance was calculated using unpaired-Wilcoxon test between sand types for each phylum (ns: $P > 0.05$; * $P \leq 0.05$; ** $P \leq 0.01$; *** $P \leq 0.001$).

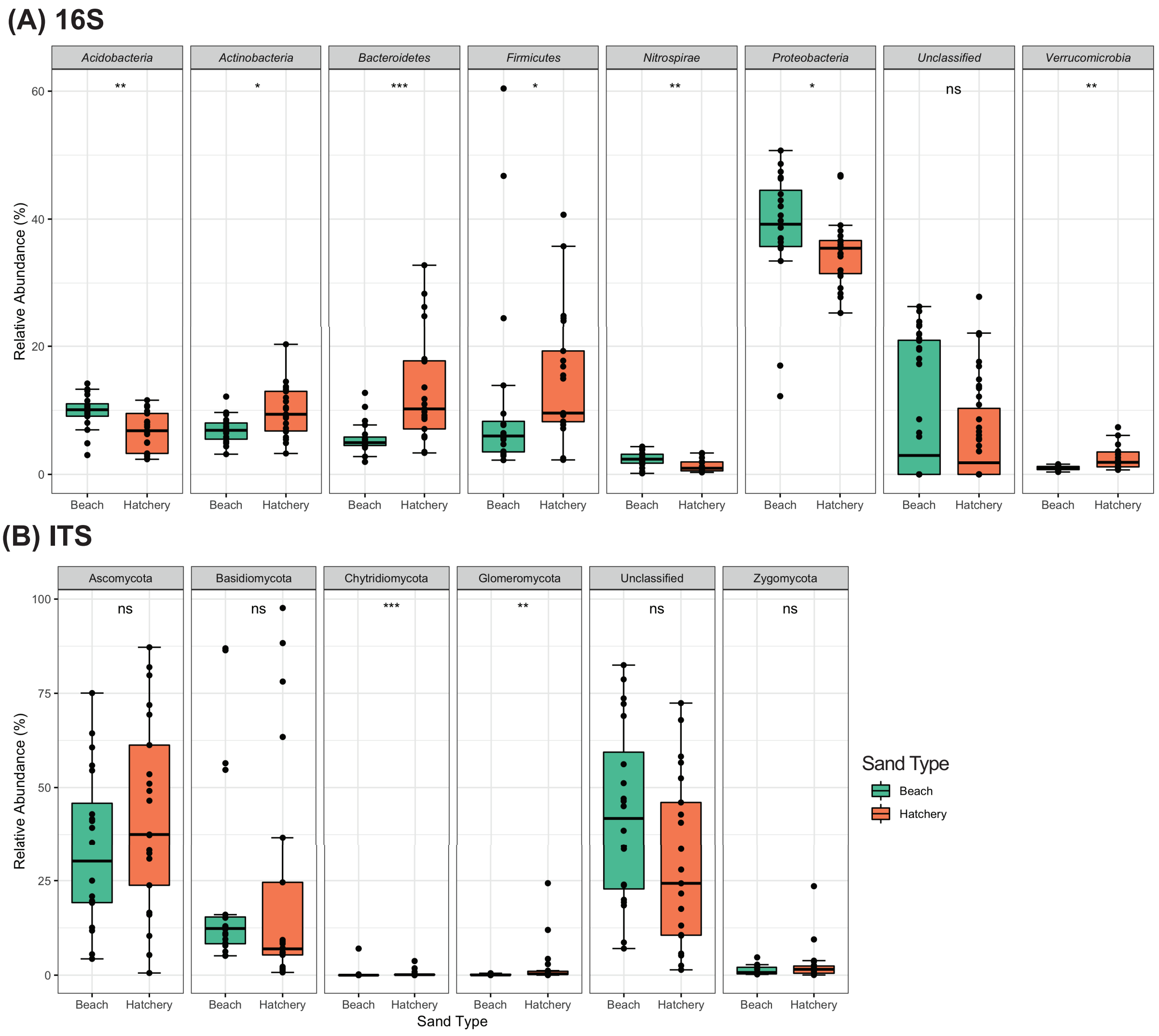

**Figure S4:** Alpha diversity measures of bacterial (A) and fungal (B) community in different sand types. Statistical significance was calculated using unpaired-Wilcoxon test between sand types for each measure (ns: $P > 0.05$; * $P \leq 0.05$; ** $P \leq 0.01$; *** $P \leq 0.001$; **** $P \leq 0.0001$).

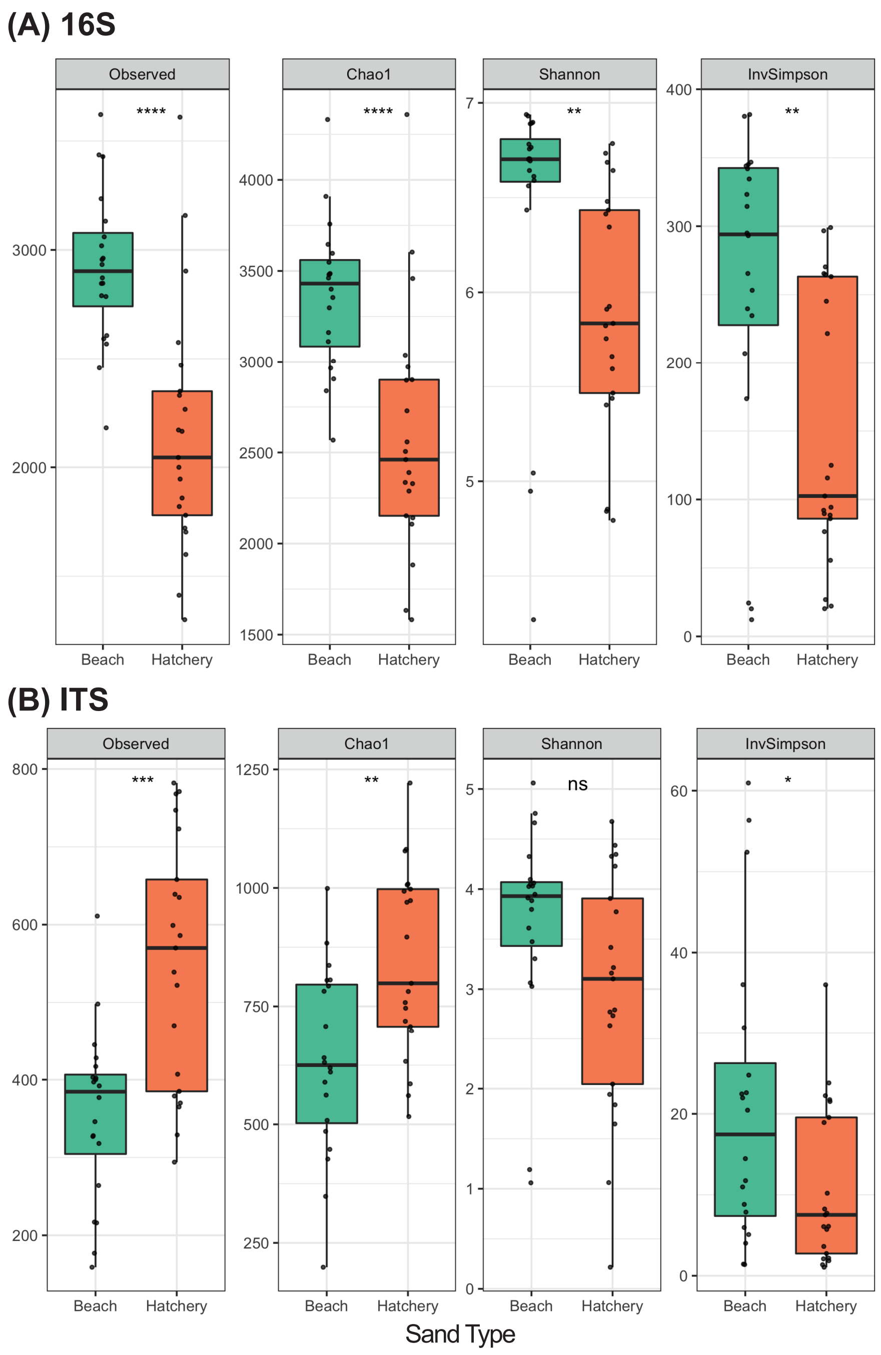

**Figure S5:** Principal Coordinate Analysis plots measured by Bray-Curtis distance of (A) bacterial and (B) fungal communities. Each dot represents one sample and different color denote sampling site. Multivariate statistical analyses ANOSIM and PERMANOVA were used to test the community differences between sand types. *Asterisk denote a subset of samples containing high relative abundance of *Inocybe* sp.

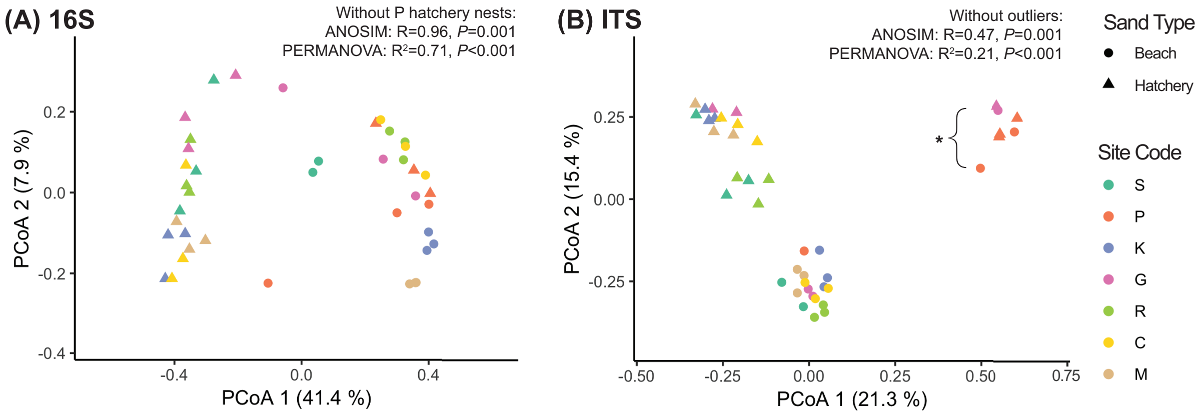

**Figure S6:** DESeq2 analysis of differentially abundant bacterial genera in different sand types. The plot shows representative taxa with $\log^2$ fold change < - 5 and > 5. Each dot denotes one genus.

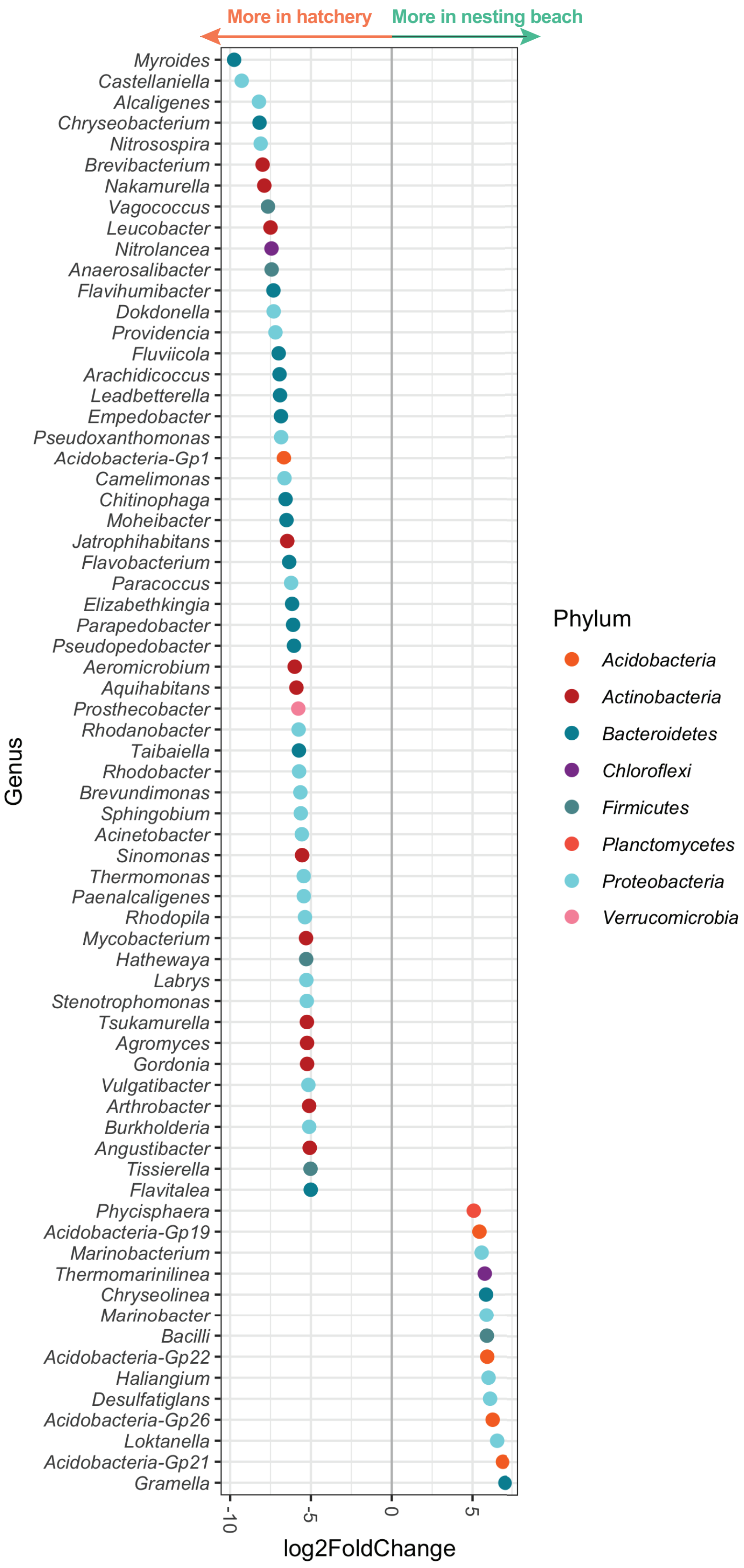

**Figure S7:** DESeq2 analysis of differentially abundant fungal OTUs in different sand types. Each dot denotes one OTU.

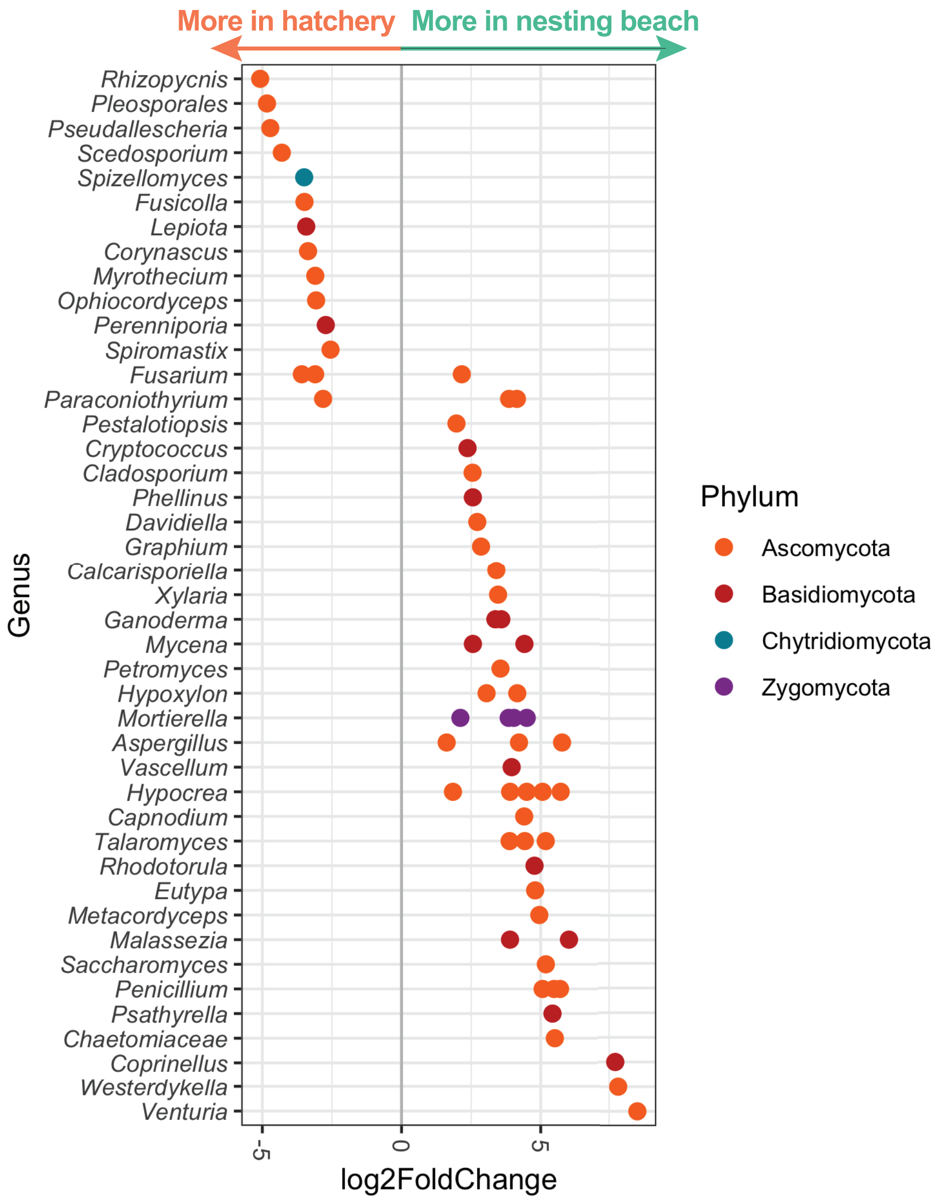

**Figure S8:** Multivariate analysis shows Principal Component Analysis biplot of different sand types with the top 14 fungal OTUs with loading scores of larger magnitudes. Description of sample label as followed: First and second alphabet in Sample ID represents site code and sand type, respectively. Samples of P hatchery sands were removed from analysis due to evidently different community structures with other hatchery sand samples.

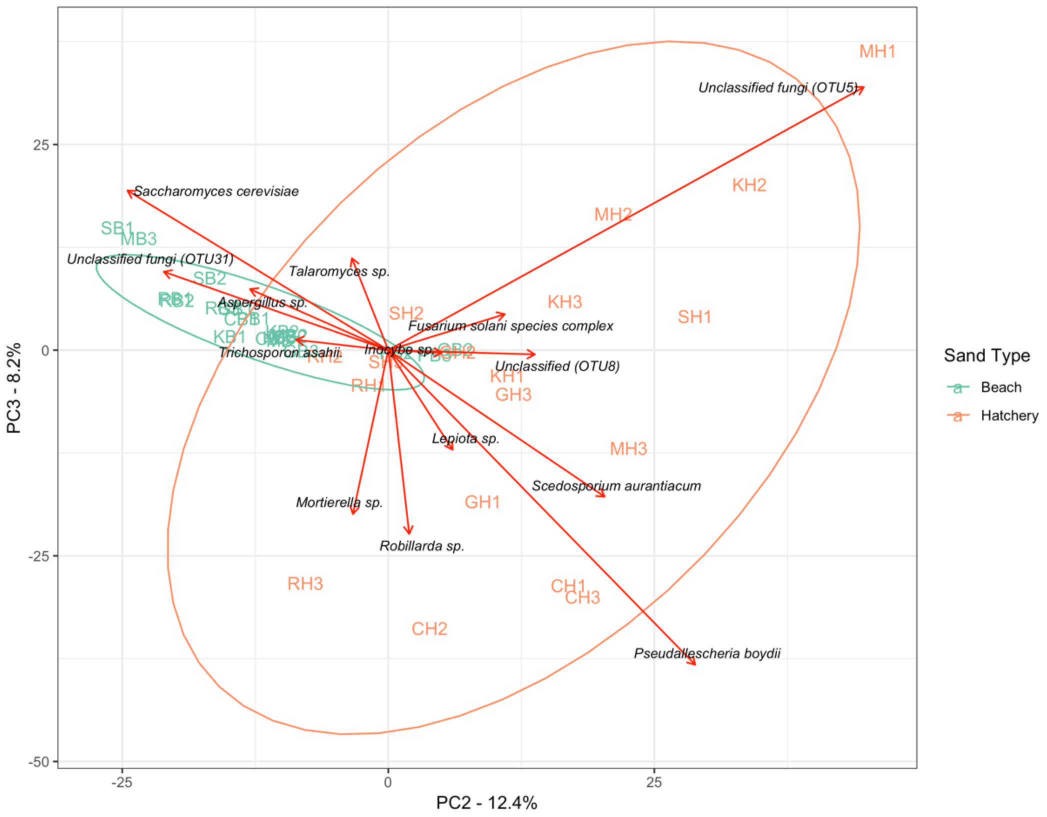

**Figure S9:** Subsets of correlation network around nodes representing *Fusarium solani* species complex, *Pseudallescheria boydii*, unclassified fungi OTU5 and their shared and direct interacting neighbors in sand of nesting beach (A) and hatchery (B). Node correspond to OTU and color represent fungi (green) and bacteria (pink). Connecting edge between nodes indicate correlations. Edge size correspond to strength of correlation. Node size in (B) correspond to number of degrees.

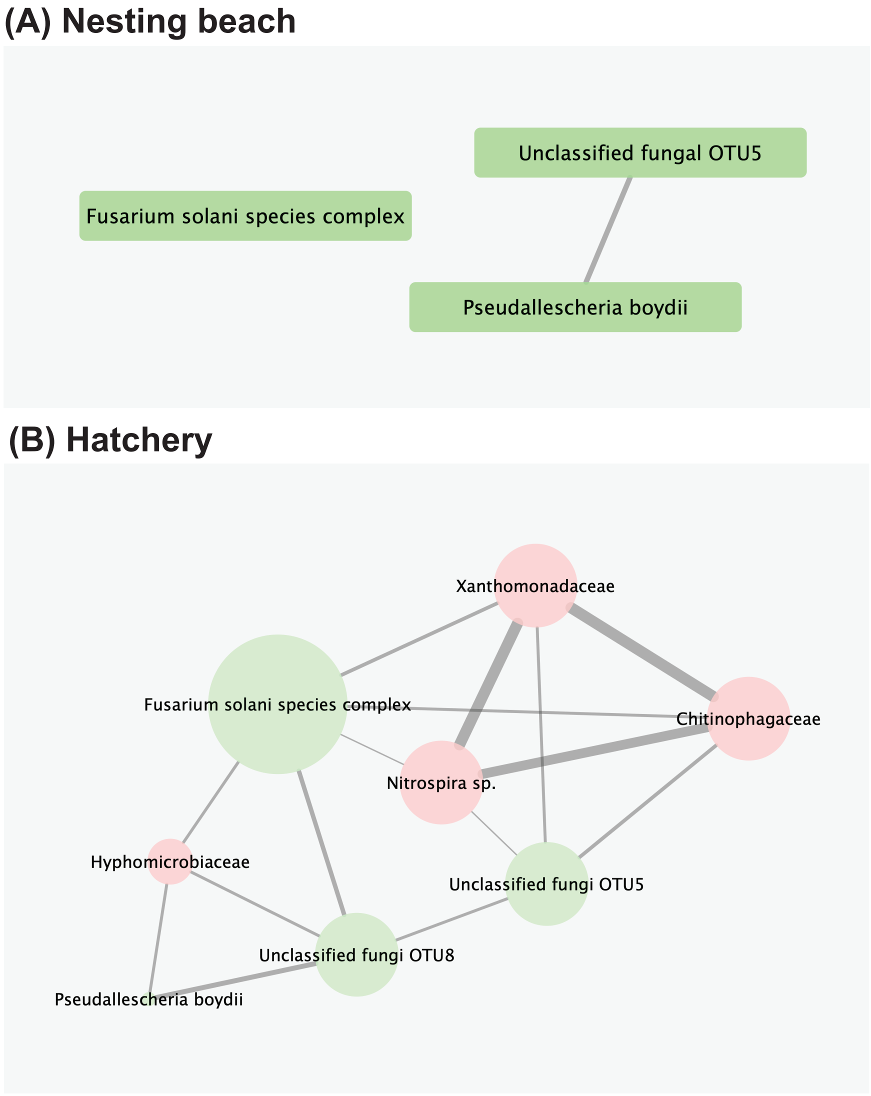
